# Spontaneous and multifaceted ATP release from astrocytes at the scale of hundreds of synapses

**DOI:** 10.1101/2022.12.05.519082

**Authors:** Yoshiki Hatashita, Zhaofa Wu, Hirotaka Fujita, Takuma Kumamoto, Jean Livet, Yulong Li, Manabu Tanifuji, Takafumi Inoue

## Abstract

Astrocytes participate in information processing by releasing neuroactive substances termed gliotransmitters, including ATP. Individual astrocytes come into contact with thousands of synapses with their ramified structure, but the spatiotemporal dynamics of ATP gliotransmission remain unclear, especially in physiological brain tissue. Using a genetically encoded fluorescent sensor, GRAB_ATP1.0_, we discovered that extracellular ATP increased locally and transiently in absence of stimuli in neuron-glia co-cultures, cortical slices, and the anesthetized mouse brain. Spontaneous ATP release events were tetrodotoxin-insensitive but suppressed by gliotoxin, fluorocitrate, and typically spread over 50–250 μm^2^ area at concentrations capable of activating purinergic receptors. Besides, most ATP events did not coincide with Ca^2+^ transients. Clustering analysis revealed that these events followed multiple distinct kinetics, and blockade of exocytosis only decreased a minor group of slow events. Overall, astrocytes spontaneously release ATP through multiple mechanisms, mainly in non-vesicular and Ca^2+^-independent manners, thus potentially regulating hundreds of synapses all together.

## Introduction

Astrocytes, the most abundant glial cells in the central nervous system, directly modulate synaptic activity and neuronal network dynamics by releasing neuroactive substances termed gliotransmitters (*1, 2*). Gliotransmission is postulated as a feedback response to neuronal activity (*3*–*5*). Astrocytes can discriminate inputs from different synaptic terminals (*6*) and utilize different gliotransmitters depending on the neuron firing frequency (*7*). These notions suggest that they can interpret surrounding neuronal activity and provide relevant feedback control. Thus, astrocytes are now regarded as partners of neurons in information processing, whereas they were classically considered to play only supportive roles in the brain.

ATP is one of the major gliotransmitters and is considered essential for brain function because impaired astrocytic ATP release leads to deficiency in synaptic plasticity (*8*), depressive-like behavior (*8*–*10*), and sleep loss in rodents (*11*). Not only is ATP released from astrocytes in response to neuronal inputs, but Koizumi et al. found tetrodotoxin-insensitive ATP release in neuron-glia co-culture (*12*). Tonic ATP release was reported in brain slice preparations (*13*–*15*), but it was unclear whether astrocytes release ATP spontaneously or in response to the basal neuronal firing.

Investigating basal ATP release activity will provide a further perspective on how astrocytes are involved in the neuron network function. Intracellular Ca^2+^ elevation in astrocytes has been implicated in ATP gliotransmission and can be triggered by various inputs including neurotransmitters, neuromodulators, changes in temperature, pH, and pressure changes, or even without stimulation (*16*–*18*). Several groups have demonstrated that optogenetically or chemogenetically evoked Ca^2+^ increase in astrocytes leads to ATP release (*19*–*21, 10*). Furthermore, spatiotemporally complex spontaneous Ca^2+^ elevations in astrocytes have been revealed (*22*–*27*), with varied areas ranging from sub-micrometer order touching a single synapse (*28*) to ones covering tens to thousands of synapses (*18*). Although the Ca^2+^-dependent gliotransmission has been the subject of debate (*29*–*32*), the above lines of evidence have prompted speculation that astrocytes locally regulate multiple synaptic activities by releasing ATP following spontaneous Ca^2+^ elevations (*18, 33*).

However, even the basic properties of ATP release, such as its spatial and temporal dynamics, remain unknown in physiological conditions due to the limitation of detection methods. Non-optical approaches using the “sniffer cell” method, which sense extracellular ligands with artificially expressed receptors specific for target molecules combined with the patch-clamp technique or Ca^2+^ imaging, and microelectrode biosensors lack spatial information to reveal the dynamics of extracellular ATP at sub-cellular resolution (*34*). Since they cannot identify individual release events, the temporal kinetics of ATP release also remains elusive. To overcome the technical difficulties, optical sensors have been developed. However, they were not adequate to detect endogenous ATP release in the brain tissue until recently (*35, 36, 34*). These limitations hampered clarification of the spatiotemporal properties of ATP release and their relationships with spontaneous Ca^2+^ activities.

We previously developed a highly sensitive genetically encoded extracellular ATP fluorescent sensor, GRAB_ATP1.0_, and successfully detected individual spontaneous ATP release events in neuron-glia culture and inflammation-induced ATP release *in vivo* (*37, 38*). In this study, we monitored extracellular ATP under physiological conditions *in vitro*, in acute brain slices, and in the mouse brain using GRAB_ATP1.0_. Furthermore, we investigated the spatiotemporal dynamics of spontaneous ATP release and its relationship with intracellular Ca^2+^ events in cortical astrocytes. Our results strongly suggest that astrocytes spontaneously and locally release ATP through multiple mechanisms, mainly in non-vesicular and Ca^2+^-independent manners.

## Results

### Spontaneous ATP release in neuro-glia co-culture is independent of neuron activity

We have previously reported that ATP release is detected without external stimulations in neuron-glia co-culture expressing GRAB_ATP1.0_ (*37*). To test whether neuronal activity participates in this spontaneous ATP signaling, we first blocked neuron firing with tetrodotoxin (TTX). The frequency and amplitude of spontaneous ATP release events were not significantly altered by the TTX treatment, while the responses were abolished by application of the P2Y1 receptor antagonist MRS-2500, as expected from the fact that the sensor originated from the human P2Y1 receptor (Fig. 1A–D). When the neuronal activity was enhanced by electrical stimulation and high K^+^ treatment, robust Ca^2+^ elevation in neurons was induced, while no increased GRAB_ATP1.0_ response was detected. The application of 100 nM ATP was sufficient to evoke a robust GRAB_ATP1.0_ response (Fig. 1E–G). Altogether, these results demonstrated that neuronal activity was not responsible for the spontaneous ATP release observed in neuron-glia co-culture.

**Fig. 1.**
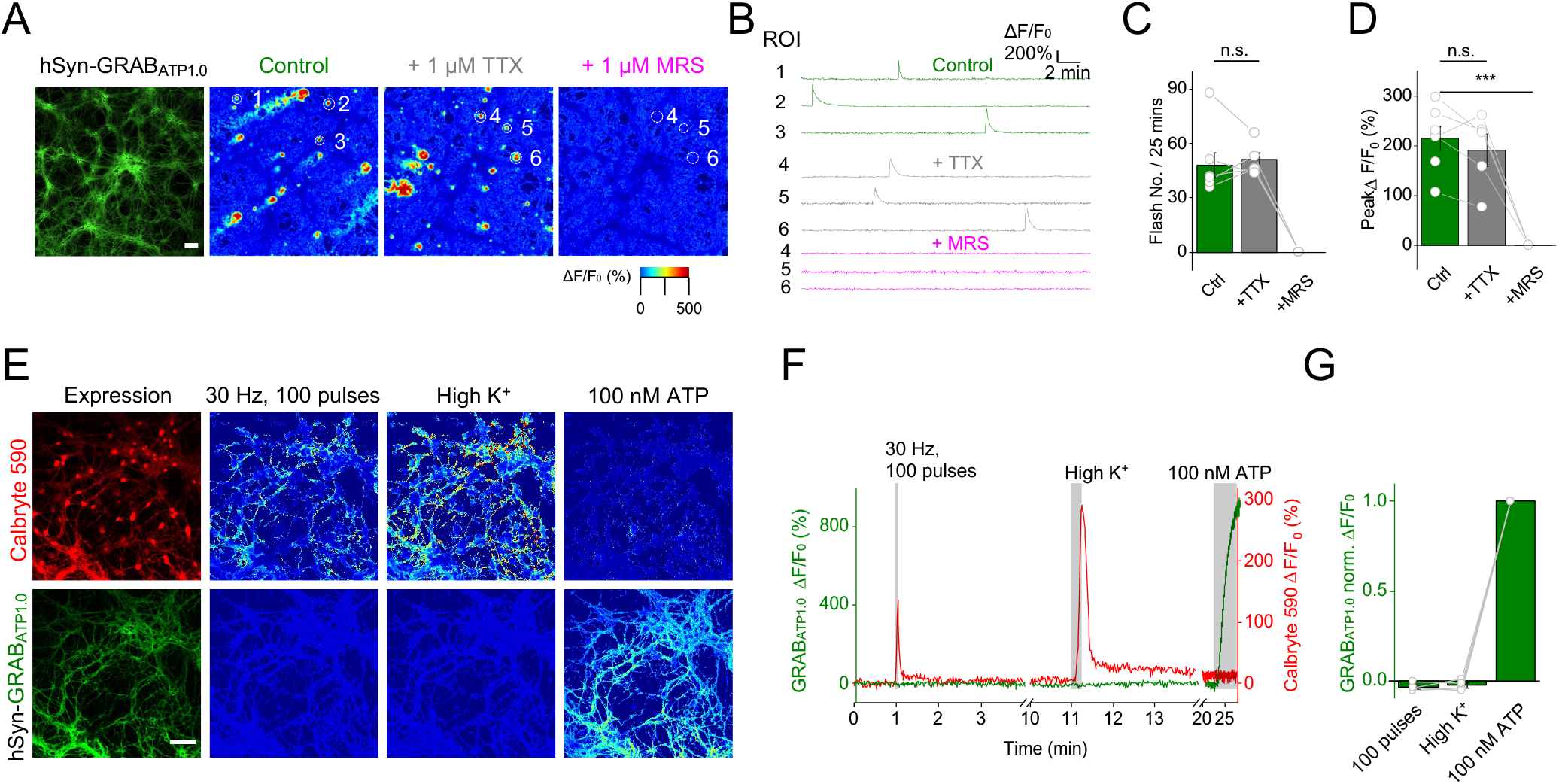
TTX-insensitive spontaneous ATP release in hippocampal neuron-glia co-culture. (**A**) Raw fluorescence image and pseudocolor images of the fluorescence response (ΔF/F_0_) of GRAB_ATP1.0_ in Tyrode’s solution (control), tetrodotoxin (TTX, 1 μM), and MRS-2500 (a P2Y1R antagonist, MRS, 1 μM). (**B**) Time course traces of the fluorescence response (ΔF/F_0_) of GRAB_ATP1.0_ in the regions of interest (ROI) circled in A. (**C–D**) Frequency (number of events per 25 min, **C**) and peak amplitude (**D**) of fluorescence transients recorded from 4–6 coverslips. (**E**) Images of Calbryte 590 (top) and GRAB_ATP1.0_ (bottom) showing raw fluorescence, ΔF/F_0_ on field electrical stimulation (100 pulses at 30 Hz), high K^+^ stimulation and 100 nM ATP application, from left to right. (**F**) Time course traces showing the mean fluorescence response (ΔF/F_0_) in the field of view of Calbryte 590 (red) and GRAB_ATP1.0_ (green) to the stimulations. (**G**) Peak amplitudes of GRAB_ATP1.0_ fluorescence responses recorded from 3 coverslips. Scale bars: 100 μm (**A**) and 30 μm (**E**). Student’s t-test, ***p<0.001; n.s., not significant.

### Cortical astrocytes spontaneously and locally release ATP

To monitor extracellular ATP dynamics in murine brain tissue, astrocytes expressing GRAB_ATP1.0_ were prepared by means of *in utero* electroporation with an improved transposon vector based on the iOn-switch (*39*) (Fig. 2A). This approach achieved sparse expression of the ATP sensor in cerebral cortex and enabled us to distinguish individual astrocytes by morphology (Fig. 2B). Neither glial scar formation nor upregulation of glial fibrillary acidic protein (GFAP) expression was observed (Fig. S1), suggesting no detectable inflammatory activation of astrocytes.

**Fig. 2.**
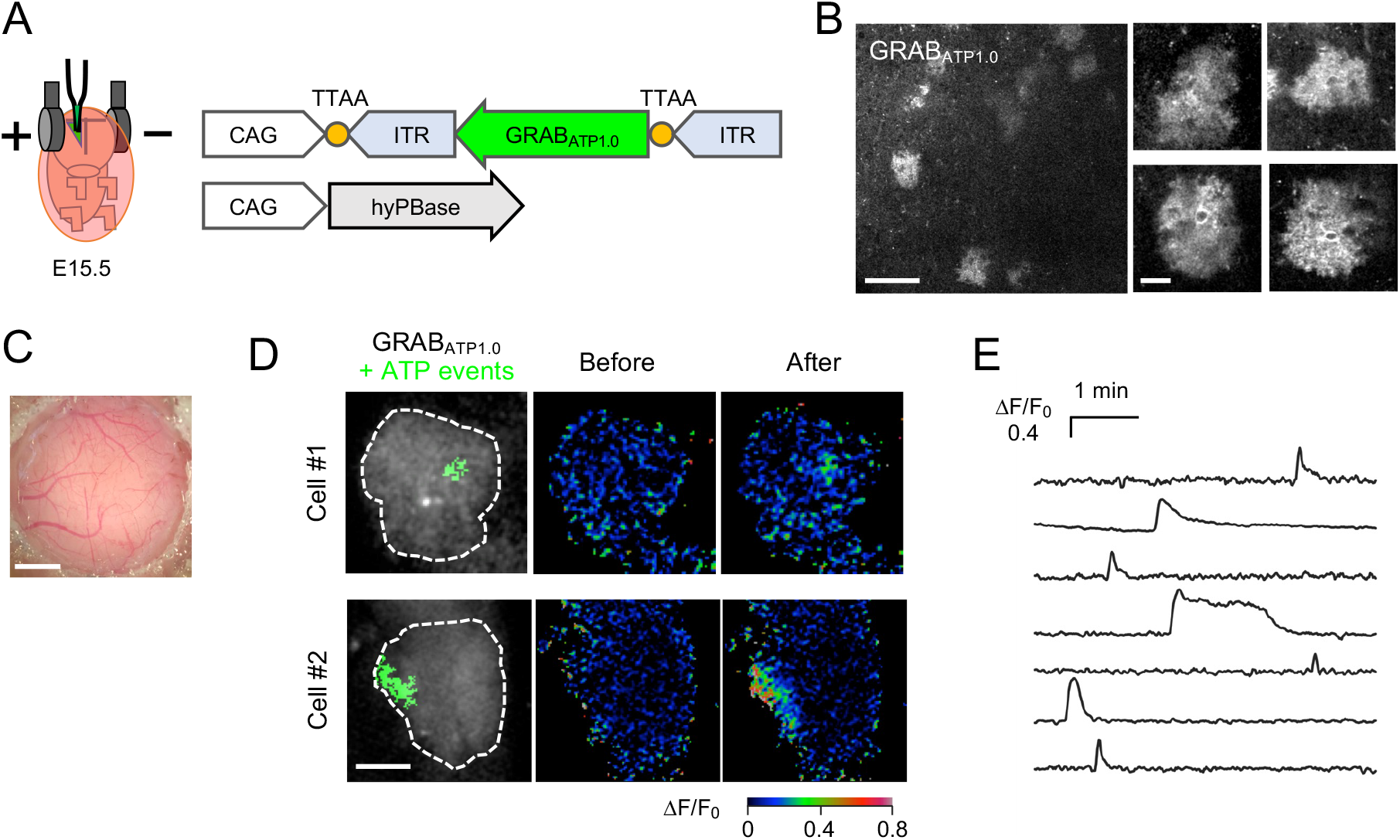
Local and transient ATP release *in vivo*. (**A**) Schematic drawing of *in utero* electroporation and plasmids for expressing GRAB_ATP1.0_ with the iOn-switch PiggyBac transposon system. (**B**) Images of cortical slices prepared from P40 mice electroporated at E15.5 showing GRAB_ATP1.0_ fluorescence expressed in a small number of astrocytes. High-magnification images (right) were taken from the same slice with different focal planes. (**C**) A thinned-skull cranial window. (**D**) Focal spontaneous ATP release events in astrocyte arbors *in vivo*. GRAB_ATP1.0_ fluorescence images of two astrocytes (left). ATP increase areas detected by AQuA software (*72*) are shown in green, and astrocyte borders are indicated by white dashed lines. ΔF/F_0_ images of GRAB_ATP1.0_ before (middle) and during (right) the spontaneous ATP events. (**E**) Time course traces of GRAB_ATP1.0_ signals (ΔF/F_0_) showing ATP release events. Scale bars: 100 μm (**B**, left), 20 μm (**B**, right, and **D**), and 1 mm (**C**).

We then monitored GRAB_ATP1.0_ fluorescence in anesthetized mice using two-photon microscopy. We found transient increases in extracellular ATP within restricted subcellular regions of astrocytes (Fig. 2C–E; area: 124.16 (37.28, 381.6) μm^2^; full width at half maximum (FWHM): 6 (5, 14) sec; peak ΔF/F_0_: 51.4% (41.7%, 65.8%); median (interquartile ranges [IQRs]) in 12 events from 30 cells in 6 mice).

Such transient and localized ATP events were also found in acute brain slices without stimulation (Fig. 3A and B). To identify the source of spontaneous ATP release in slices, the activities of neurons and of astrocytes were suppressed by TTX and an astrocyte-specific gliotoxin, fluorocitrate (FC), respectively. ATP release events were insensitive to TTX concerning their frequency, area, duration, and peak amplitude (Fig. 3C–H), consistent with the results in neuron-glia co-cultures. By contrast, the frequency of ATP release was suppressed by FC (Fig. 3I and J), indicating that the spontaneous ATP release in the brain slice was dependent upon the astrocyte activity. These results indicate that cortical astrocytes release ATP independently of neuronal activity.

**Fig. 3.**
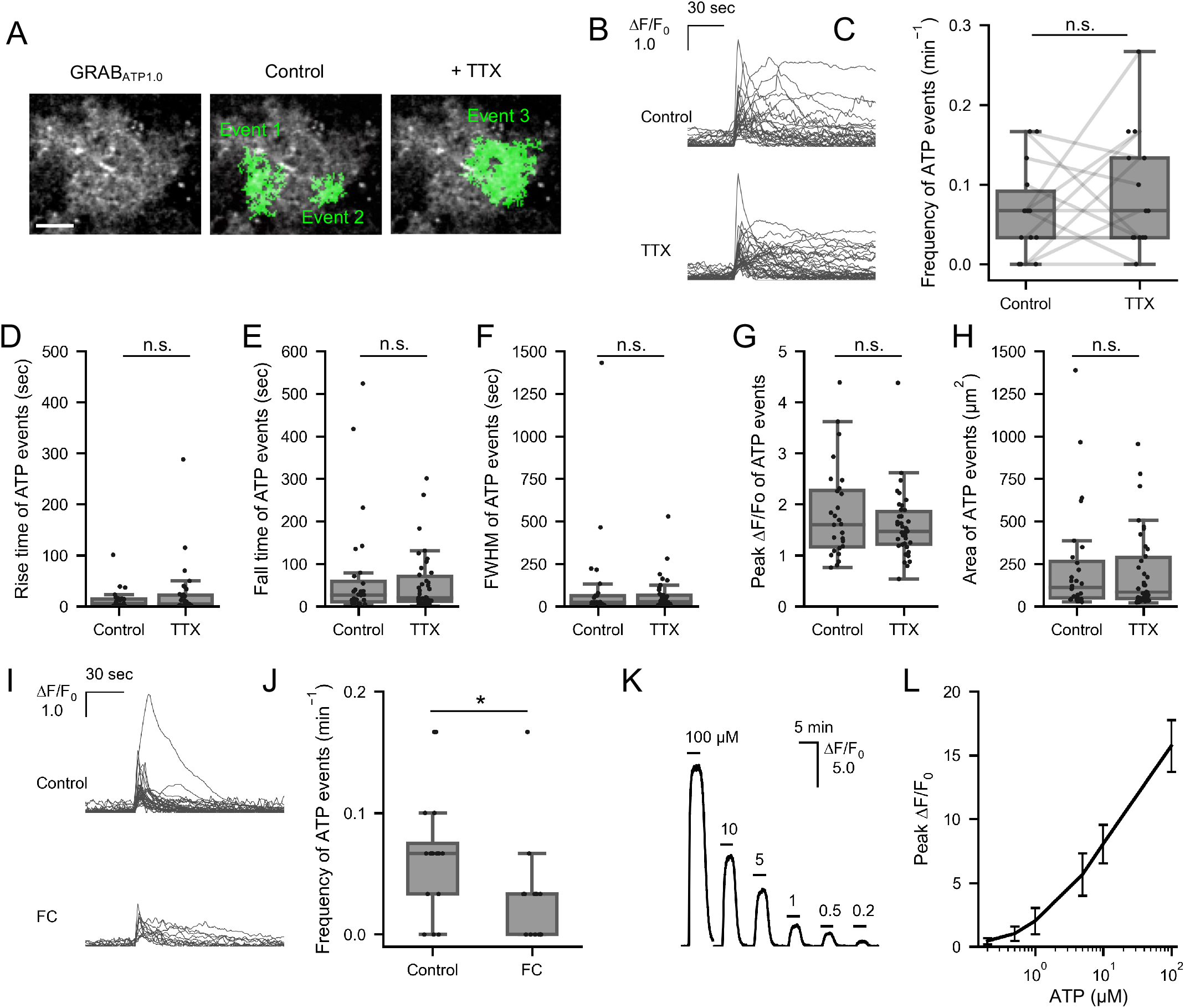
Astrocytes spontaneously release ATP in acute slices. **(A)** A cortical astrocyte in an acute slice expressing GRAB_ATP1.0_ (left) overlayed with focal spontaneous ATP release events (green) observed in control (middle) and in 1 μM TTX (right) conditions. Scale bar: 20 μm. (**B**) Time course changes in GRAB_ATP1.0_ signal (ΔF/F_0_) of spontaneous ATP release events detected in control (top) and in TTX (bottom) conditions. Events are aligned by the initial rise timing (30% of the peak amplitude). (**C**) Frequency of spontaneous ATP release events (n = 14 cells each). (**D–H**) Rise time (**D**), fall time (**E**), full width at half maximum (FWHM) of duration (**F**), peak amplitude of ΔF/F_0_ (**G**), and area (**H**) of spontaneous ATP release events in control (n = 28 events) and TTX (n = 38 events) groups. (**I**) Time course changes in GRAB_ATP1.0_ signal (ΔF/F_0_) of spontaneous ATP release events detected in control and fluorocitrate (FC, 100 μM) conditions. (**J**) Frequency of spontaneous ATP release events in the control (n = 16 cells) and FC (n = 15 cells) groups. (**K**) ΔF/F_0_ changes of GRAB_ATP1.0_ expressed in an astrocyte in acute slice responding to bath-application of 100, 10, 5, 1, 0.5, and 0.2 μM ATP. (**L**) Calibration of the response of GRAB_ATP1.0_ expressed in astrocytes in acute slice (mean ± S.D, n = 5 cells). Wilcoxon signed-rank test (**C**) and Mann-Whitney U test (**D–H** and **J**) were used. Box plots show the median, 25th, and 75th percentile.

To estimate the concentration of locally released ATP, the GRAB_ATP1.0_ response in slice was calibrated by bath-application of ATP (Fig. 3K and L). With this calibration result and the peak amplitudes of ATP release events (Fig. 3G), the peak concentration of typical ATP release appeared to range between 0.5–5 μM, which meets the concentration range for activation of purinergic receptors (*40, 41*), indicating that spontaneous ATP release may affect multiple synapses within the local areas all at once.

### The majority of spontaneous ATP release do not coincide with Ca^2+^ elevation in astrocytes

To examine the relationship between spontaneous ATP release and Ca^2+^ activity in astrocytes, we monitored extracellular ATP and intracellular Ca^2+^ dynamics simultaneously by expressing GRAB_ATP1.0_ and a membrane-anchored red Ca^2+^ sensor, Lck-REX-GECO1 (*42*), in astrocytes in acute brain slices (Fig. 4A). Of note, the frequency of Ca^2+^ transients did not increase over time during the 30-minute recording session (Fig. S2), ruling out the possibility of photo-damage caused by two-photon excitation (*43*). The rise time, fall time, FWHM of duration, and area of ATP events were relatively more widely distributed than those of Ca^2+^ events (Fig. 4B–E). In contrast, the peak amplitude and subcellular occupancy of ATP and Ca^2+^ events were distributed similarly (Fig. 4F and G). Unexpectedly, more than half of the spontaneous ATP release events did not coincide with Ca^2+^ elevation (Fig. 4K). ATP release events devoid of Ca^2+^ events were also detected through line-scanning at a 100 Hz sampling rate (Fig. S4). Among the spatially overlapping patterns, ATP events followed by Ca^2+^ elevation (Fig. 4H) were most frequently observed, and the incidences of ATP events following or coinciding with Ca^2+^ elevation were low (Fig. 4I and J). The occurrence of ATP events preceding or following Ca^2+^ events was not statistically significant, although that of the coinciding events was above chance-level (p < 0.001; Fig. 4K). Besides, Ca^2+^ events occurred with significantly higher frequency only during the 30-second period just after the ATP release events (Fig. 4L). The frequency of Ca^2+^ events per cell did not correlate with that of spontaneous ATP events (Fig. 4M). These results suggest that most of the spontaneous ATP release occurred in a Ca^2+^-independent manner.

**Fig. 4.**
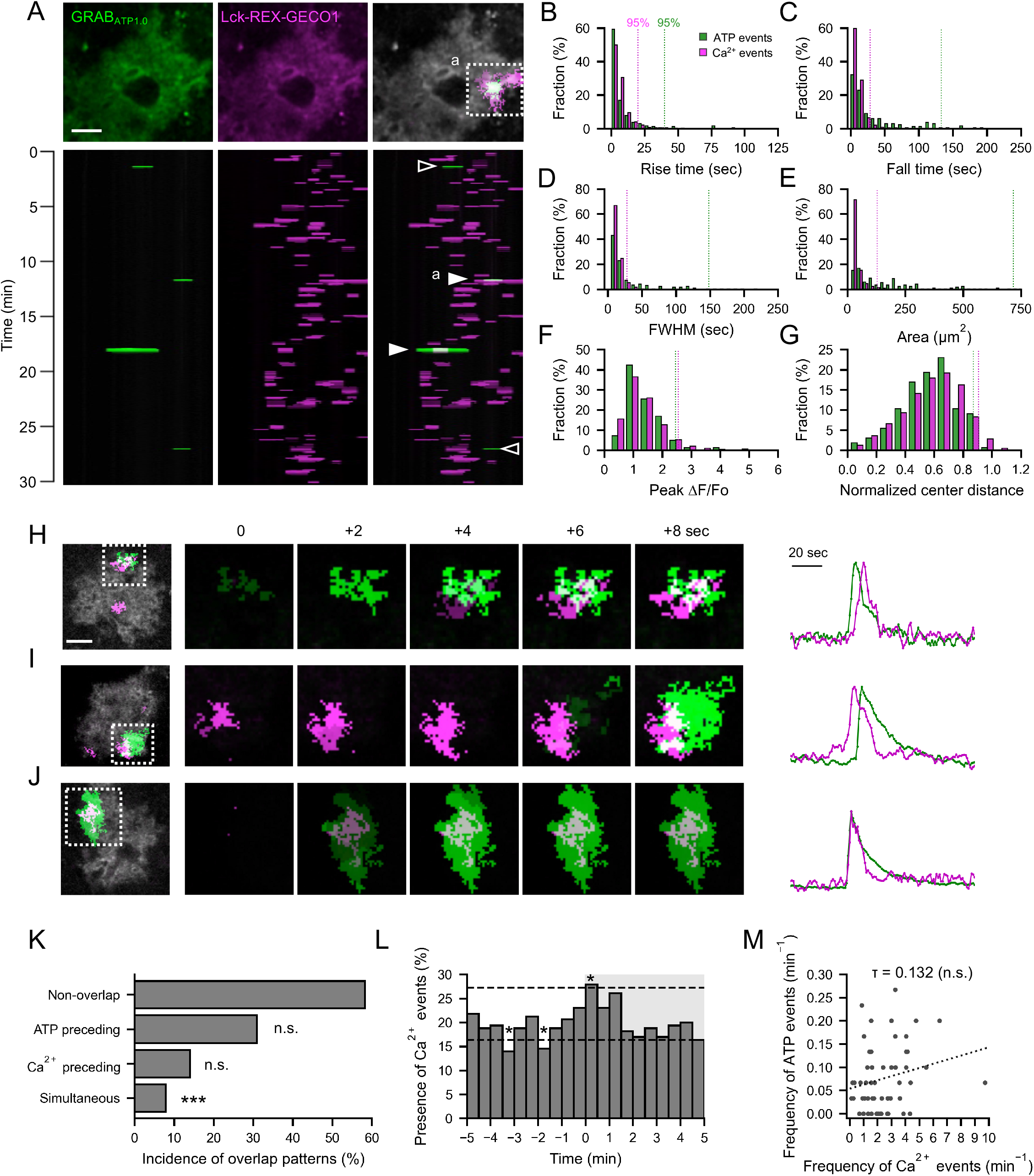
Simultaneous imaging of spontaneous ATP release and Ca^2+^ activity in astrocytes in acute slices. (**A**) X-T binary maps showing overlays (maximal values on the y-axis) of spontaneous ATP release events (bottom left), spontaneous Ca^2+^ events (bottom, middle), and the both (bottom right) detected from an astrocyte expressing GRAB_ATP1.0_ (top left) and Lck-REX-GECO1 (top middle). ATP events occurred with (close arrowhead; X-Y images of the both events (**a**) are overlaid in the top right image) or without (open arrowhead) accompanying Ca^2+^ events at the same time. (**B–G**) Histograms showing the distribution of rise time (**B**), fall time (**C**), FWHM of duration (**D**), area (**E**), peak amplitude in ΔF/F_0_ (**F**), and distance between the gravity center of event area and the gravity center of astrocyte area normalized by the cell radius (**G**) of spontaneous Ca^2+^ transients (magenta, n = 3663 events) and ATP release events (green, n = 165 events). Dashed lines indicate the upper boundaries of 0–95% density. The numbers of events out of the abscissa range are 2, 2, 2, 8, 0, and 0 in the ATP events and 19, 4, 1, 17, 1, and 3 in the Ca^2+^ events in **B**–**G**, respectively. (**H–J**) Three representative data showing overlapping ATP release and Ca^2+^ events (top, an ATP event preceding a Ca^2+^ event; middle, an ATP event following a Ca^2+^ event; bottom, simultaneous events) in sequential images (left) and time course traces of ΔF/F_0_ (right). (**K**) Incidences of spontaneous ATP release events classified by temporal overlapping patterns with Ca^2+^ events. Chance levels (***p < 0.001; n.s., not significant) were estimated by shuffling the onset time of Ca^2+^ events (10,000 iterations). (**L**) The proportion of ATP release events in which Ca^2+^ events were observed in 30-second time windows. The onset of ATP release events corresponds to time 0. Chance levels (dashed lines, 95%) were estimated by shuffling the onset time of Ca^2+^ events (10,000 iterations). (**M**) Correlation between the frequency of spontaneous ATP release events and that of Ca^2+^ transients (Kendall’s rank correlation, n = 55 cells). Scale bars: 20 μm.

### ATP release events can be classified into five clusters based on the waveform

Vesicular release and release through several channels have been proposed as candidate mechanisms for ATP release from astrocytes (*44*). Because spontaneous ATP release events appeared to vary kinetically (Fig. 3B and 4B–E), we presumed that their waveforms might reflect distinct ATP release mechanisms. To classify ATP events according to their waveforms, hierarchical clustering was conducted using a shape-based distance (SBD) algorithm based on normalized cross-correlation (*45*) (Fig. S5). Two classes and five subdivided clusters were determined, and ATP events in the primary class (class 2) had sharper waveforms than the other class (Fig. 5A and B). Additionally, non-linear dimensionality reduction by the uniform manifold approximation and projection (UMAP) algorithm (*46*) clearly showed gaps among the clusters determined by hierarchical clustering, except for clusters iii (brown) and v (purple; Fig. 5J). Altogether, these analyses revealed that the spontaneous ATP release from astrocytes comprises multiple groups presenting distinct kinetics.

**Fig. 5.**
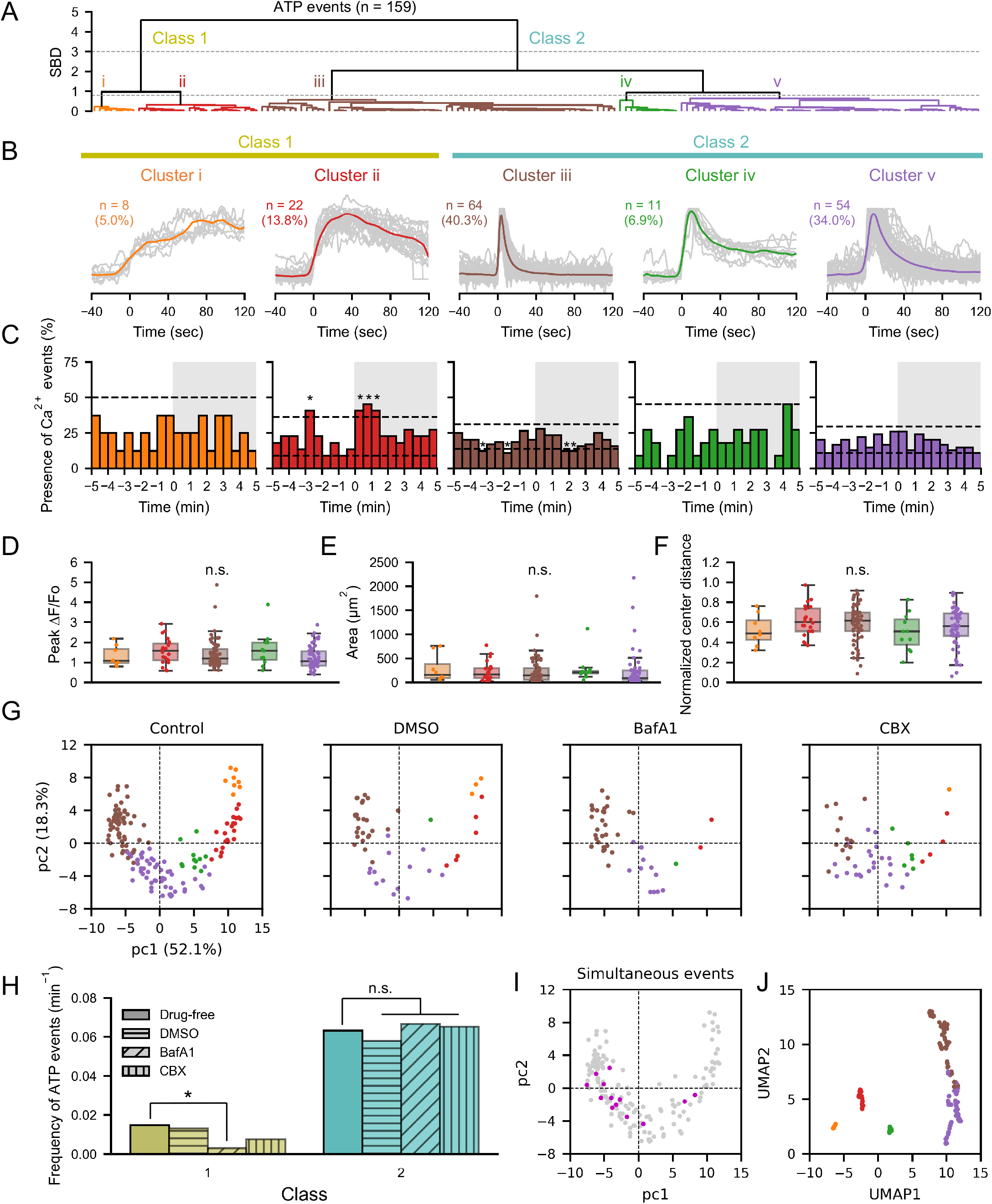
Clustering analysis of spontaneous ATP release events based on waveform. **(A)** Spontaneous ATP release events were classified into two classes and five sub-divided clusters by hierarchical clustering based on the waveform. (**B**) Gray lines indicate traces normalized by the peak amplitude of ΔF/F_0_, and colored lines represent the barycenter of traces belonging to the cluster. (**C**) The proportion of ATP release events in which Ca^2+^ events were observed in 30-second time windows. The onset of ATP release events corresponds to time 0. Chance levels (dashed lines, 95%) were estimated by shuffling the onset time of Ca^2+^ events (10,000 iterations). (**D–F**) The peak amplitude of ΔF/F_0_ (**D**), area (**E**), and distance between the gravity center of event area to the gravity center of cell area normalized by the cell radius of ATP release events in each cluster (median, and 25th and 75th percentile; Mann-Whitney U test). (**G**) Distribution of ATP release event waveforms in 2D by reducing the dimensions with the principal component analysis. The first and second principal components (pc1 and pc2) indicate 52.1% and 18.3% of explained variance, respectively. Colors correspond to those of the clusters in **A**. (**H**) Effects of inhibitors on the ATP release event frequency in the two classes (control: n = 68 cells; DMSO: n = 23 cells; BafA1 (2 μM): n = 22 cells; Carbenoxolone (CBX, 100 μM): n = 22 cells; statistics: Poisson distribution). (**I**) ATP release events occurring simultaneously with Ca^2+^ transients (13 events) are colored magenta, and others gray in the same scatter plot shown in **G** (Control). (**J**) Distribution of the ATP release event waveforms displayed by reducing the dimensionality using the uniform manifold approximation and projection (UMAP) method. *p < 0.05; n.s., not significant.

We then examined the relationship between ATP and Ca^2+^ events in each cluster. We found that only the ATP events in cluster ii tended to accompany Ca^2+^ events before and just after the ATP events (Fig. 5 C), which was buried in the overall analysis (Fig. 4L). This implies that ATP event types classified by the waveform reflect different mechanisms that lead to different ATP kinetics. No apparent difference was found in peak amplitude and area of ATP events among the clusters (Fig. 5D and E), suggesting that the amount of released ATP may be irrelevant to the distinct effects of ATP clusters on the Ca^2+^ events. Also, preference for the subcellular occupancy of ATP release was not observed among the clusters (Fig. 5F).

### A minority of ATP events characterized by slow kinetics correspond to vesicle release

To verify the possibility that the difference in the waveform of ATP events reflects distinct release mechanisms, ATP events observed in the presence of inhibitors against vesicular release (bafilomycin A1, BafA1) or hemichannels (carbenoxolone, CBX) were similarly classified using the nearest neighbor classification algorithms with the SBD measure. Hemichannels were reported to participate in ATP release in the resting state of hippocampal slices (*13*). Notably, the frequency of ATP events belonging to class 1 decreased under BafA1 treatment, but not in the DMSO-treated and control groups (Fig. 5G and H). Also, the BafA1 treatment abolished ATP events assigned to cluster i (Fig. S6). These results suggest that the class 1 ATP release events characterized by a slow time course are driven by vesicular release. In contrast, treatment with CBX showed no apparent changes in any of the classes or clusters, suggesting that the contribution of connexin and pannexin hemichannels is minimal in cortical slices (Fig. 5G and H, and Fig. S6). Note that the ATP release events that simultaneously occurred with Ca^2+^ events mostly belonged to class 2 (Fig. 5I; 12 out of 13 simultaneous events), implying that they correspond to channels permeable to both ATP and Ca^2+^, or rely on Ca^2+^-dependent ATP channel opening, though their occurrence was low (12 out of 129 ATP events in class 2).

## Discussion

In this study, we monitored spontaneous extracellular ATP dynamics using GRAB_ATP1.0_ in neuron-glia co-culture, acute brain slices, and live brain. With the high sensitivity and spatiotemporal resolution of the sensor, ATP release events were spatiotemporally detected without stimuli in the three preparations. We then investigated the mechanisms of the spontaneous ATP release in acute slice and obtained three major findings: first, astrocytes release ATP independently of nearby neuronal activities; second, dual-color imaging revealed that ATP release events rarely coincide with a spontaneous Ca^2+^ transient and do not correlate with the astrocytic Ca^2+^ event frequency; and finally, clustering analysis suggested that the spontaneous ATP release events involve multiple mechanisms with non-vesicular release being the major one, and that long-lasting ATP release is driven by exocytosis.

### Astrocyte actively drives purinergic signaling

Tonic extracellular purinergic signaling deriving from astrocytes in unstimulated brain slices has been reported (*13, 14*), but whether astrocytes release ATP in a spontaneous fashion or as a consequence of basal neuronal activity has not been elucidated. In the present study, we demonstrated that astrocytes release ATP independently of neuron activity (Fig. 3A–J), prompting the notion that astrocytes can provoke active communications via gliotransmission spontaneously in addition to their classical role as passive elements of neuronal processing.

### Spatial properties of ATP release from astrocytes

Despite the morphological hallmarks where individual astrocytes interact with tens of thousands of synapses within their arbor territories, the spatiotemporal properties of gliotransmitter release have not been determined due to the limitations of detection methods. Our extracellular ATP imaging approach using GRAB_ATP1.0_ revealed that spontaneous ATP release events typically spread over 50–250 μm^2^ at a concentration of 0.5–5 μM in the slice. Considering the synapse density in the murine cortex (∼0.7 synapses/μm^3^; (*33, 47*)) and EC_50_ of purinergic receptors (i.e., P2X1: 0.07; P2X2: 1.2; P2X3: 0.5; P2X4: 10; P2Y1: 8 μM; (*40, 41*)), our results indicate that individual ATP release events can potentially affect hundreds of nearby synapses all together even in a single focal plane. Halassa et al. speculated that gliotransmission groups adjacent synapses into functional ensembles because Ca^2+^ elevation that triggers gliotransmitter release is confined to subcellular regions in astrocytes (*33*). The present results on ATP show that most of the release events of this particular gliotransmitter do not accompany Ca^2+^ events but that their spread is indeed localized. Thus, the conclusion by Halassa et al. that adjacent synapses form functional groups by local gliotransmission sounds valid. It should be noted, however, that GRAB_ATP1.0_ captures both ATP and its degradation product, ADP (*37*). Therefore, the spatial and temporal spread of ATP, not ADP, could be smaller than that estimated in this study.

### ATP-induced Ca^2+^ response in astrocytes

It is reported that extracellularly released ATP induces astrocytic Ca^2+^ propagation in culture (*12, 48*) and in pathological conditions *in vivo* (*49*). It evokes an astrocytic Ca^2+^ response in an autocrine fashion in the brain slice (*19*). Our result showing that the frequency of Ca^2+^ activity slightly increased just after ATP release (Fig. 4L) may reflect such ATP-induced Ca^2+^ response, though only in a quarter of the entire ATP release events observed, suggesting that spontaneously released ATP has little impact on astrocytic Ca^2+^ dynamics in the physiological slice condition. Our clustering analysis suggests that the spontaneous ATP gliotransmission involves multiple types of release machinery, and cluster ii only showed an enhancement of Ca^2+^ activity after ATP release (Fig. 5C), which led us to presume that distinct ATP release mechanisms govern different downstream effects and also cooperatively engage in basal purinergic signaling.

### Mechanisms of spontaneous ATP release from astrocytes

Our pharmacological results suggest that ATP release events of relatively slow kinetics (class 1) are driven by vesicular ATP release. The slower decay phase of class 1 events could be explained by slower ATP diffusion, slower ATP degradation by ectonucleotidase, and long-lasting ATP release. Since it is hard to imagine that the ATP diffusion differs from its degradation conditions much in the extracellular space long-lasting ATP release sounds reasonable. The asynchronous vesicle fusion mechanism, which does not accompany an active zone-like structure or a pool of vesicles (*50*), fits with such a release pattern. Studies directly monitoring vesicular ATP release in astrocyte culture reported several minutes lags between evoked Ca^2+^ elevation and resultant exocytosis. Of note, the occurrence of Ca^2+^ events in cluster ii of class 1 was high during the 30-second period three minutes before the ATP events (Fig. 5B), which may indicate that cluster ii is driven by Ca^2+^-triggered exocytosis. Then, given that most of the Ca^2+^ events were not accompanied by ATP release (Fig. 4A), a yet-known triggering mechanism for such asynchronous vesicular ATP release other than the Ca^2+^ signal can be presumed. Alternatively, specific spatiotemporal patterns and/or strength of the Ca^2+^ signal could be effective. Besides, cluster i in class 1 is probably attributed to Ca^2+^-independent exocytosis, which is also reported in the study of astrocyte culture (*51*). Thus, the mechanisms evoking vesicular ATP release from astrocytes remain an open question.

On the assumption that the class 2 ATP release identified in this study represents channel-mediated ATP release, we can infer that this mechanism predominates over exocytosis (class 1; Fig. 5A) in the basal purinergic signaling. It has been postulated that channel-mediated ATP release engages under pathological conditions (*52, 53*). However, accumulating evidence suggests that these mechanisms are also involved in basal ATP gliotransmission (*54, 55, 13, 8, 15*). While connexin hemichannels (*13*), pannexin 1 (*56, 55*), calcium homeostasis modulator 2 (*8*), volume-regulated anion channels (*57*), and maxi-anion channels (*58*) have been identified as ATP release channels in astrocytes, results in this study with CBX (Fig. 5H) indicate that connexin hemichannels and pannexin 1 are less plausible candidates for the spontaneous ATP release events in the cortex. We assume that the primary ATP release channel in the cortex is one of the anion-selective channels because of the very low incidence of synchronous Ca^2+^ and ATP events. To date, however, the molecular identity of these anion channels has not been elucidated, and there are no specific blockers. Further studies on ATP release channels are needed to understand the molecular mechanisms underlying the spontaneous ATP release from astrocytes.

Our results showing that intracellular Ca^2+^ elevation is less likely responsible for the spontaneous ATP release events except for those in cluster ii, indicate that Ca^2+^-independent processes may play crucial roles in the basal purinergic signaling. Various environmental changes such as decrease and increase in extracellular Ca^2+^ and K^+^ concentrations, respectively, cell swelling, low oxygen, and mechanical transduction have been demonstrated to open ATP release channels (*59*), while it is not clear whether actual changes in those parameters are sufficient to affect ATP release channels under physiological conditions. Besides, intracellular Ca^2+^ events could still be candidates for ATP release: faint nanoscale Ca^2+^ events in the astrocyte structure only detectable by STED microscopy (*28*) or out-of-focus Ca^2+^ events might have been missed in this study. Thus, future studies employing advanced optical techniques including super-resolution microscopy and volumetric multi-photon imaging might help uncover the mechanisms underlying the spontaneous ATP release from astrocytes in space and time.

### Physiological significance of spontaneous ATP release

To date, few reports have been published on the physiological significance of the spontaneous ATP release from astrocytes. Pioneering studies on spontaneous ATP release in neuron-glia co-culture reported that extracellular ATP suppresses the excitatory neuron activity through presynaptic P2Y receptors, and 0.3–3 μM of extracellularly applied ATP was sufficient to attenuate synaptic release (*12, 60*). As for evoked-ATP release, ATP upregulates the activity of CCK-positive interneuron via presynaptic P2Y1 receptor while downregulating pyramidal neuron activity mediated by its metabolite adenosine (*61*), and it attenuates inhibitory postsynaptic currents via postsynaptic P2X2 and P2X4 receptors (*62*) in the slice. In these evoked-ATP release studies, an increase in extracellular ATP was estimated at about 1 μM; therefore, we expect that the spontaneous ATP release observed in this study potentially leads to similar effects on neurons. Moreover, it is also plausible that the spontaneously released ATP modulates not only the neuron activity but also the microglial surveillance or the blood flow control, which are associated with purinergic signaling (*63, 64*). Besides, as the loss of ATP release from astrocytes is related to psychiatric symptoms (*8*–*10*) and pathological conditions promote hyperactive extracellular ATP signaling (*49, 37, 44*), future studies revealing how the spontaneous purinergic signaling of astrocytes is altered in pathological conditions will provide further insights into the roles of astrocytes both in physiological and pathophysiological conditions.

### Significance

The results presented in this study indicate that astrocytes spontaneously release ATP through multiple mechanisms, mainly in non-vesicular and Ca^2+^-independent manners, which potentially coordinate hundreds of synapses at once. Our study, thus, provides three insights into astrocyte-neuron interaction mediated by gliotransmitter release: first, contrary to the classical regard for astrocytes as a passive element in the brain, astrocytes can provoke active communications independently of neuronal activity; second, our findings further strengthen the concept that neighboring synapses form functional islands regulated by purinergic signaling derived from astrocytes; third, although Ca^2+^ activity is considered vital for astrocyte function, Ca^2+^-independent processes may also play crucial roles in the gliotransmitter release.

### Limitations of the study

The present study has disclosed the spontaneous and localized release of ATP from astrocytes in culture and brain tissue, where we explored the mechanisms of this release, especially in slice preparation. Anesthetized mice, where neuronal inputs are depressed, also exhibit transient ATP release, however, its detailed mechanisms need to be thoroughly examined *in vivo*. Besides, while our work shed light on the spontaneously derived-ATP signaling in the cortex, further studies exploring other gliotransmitter dynamics in various brain regions are awaited to fully understand astrocyte-mediated active communications.

## Materials and Methods

### Experimental Design

All procedures were approved by the Committee on the Ethics of Animal Experiments of Waseda University and the Animal Care and Use Committees at Peking University. Wild-type ICR mice were maintained in Waseda University in accordance with the guidelines outlined by the Institutional Animal Care and Use Committee of Waseda University, and both male and female mice were used for experiments.

### DNA plasmid construction

Plasmid vectors based on the integration-coupled on (iOn) gene expression switch system conditioning transgene expression to piggyBac-mediated transposition (*39*) were employed for stable expression of fluorescent sensors in astrocytes. To construct the ^*iOn*^*CAG∞GRAB*_*ATP1.0*_ plasmid, the GRAB_ATP1.0_ coding sequence was amplified by PCR (primers: 5’; GTGGCCACTCGAGGATCCACCATGGAGAGAGACAC, 3’; GCACTAAAGTCGGATCAGAGGGGCGCTAGCTTACA) and cloned into ^*iOn*^*CAG ∞MCS* at the XhoI site by SLiCE cloning (*65*). ^*iOn*^*CAG∞REX-GECO1* was constructed by replacing the sequence between the BamHI and NheI sites of ^*iOn*^*CAG ∞GRAB*_*ATP1.0*_ with the BamHI-XbaI fragment of *pCMV-REX-GECO1*. For the construction of ^*iOn*^*CAG∞Lck-REX-GECO1*, the Lck domain digested from *pCMV-Lck-MaLionR* with NheI and BamHI was inserted into the AvrII and BamHI sites of ^*iOn*^*CAG∞REX-GECO1*.

### *In utero* electroporation

Pregnant ICR mice were purchased from Japan SLC Inc., and *in utero* electroporation (IUE) was performed as previously described (*66*) with the following modifications. Mice at gestation day 15 were deeply anesthetized with either a mixture of medetomidine (0.3 mg/kg body weight), midazolam (4 mg/kg) and butorphanol (5 mg/kg) or with sodium pentobarbital (50 mg/kg) by intraperitoneal administration. Approximately 1 μL of plasmid solution containing 2.0 μg/μL of the iOn-switch vectors, 0.5 μg/μL of *pCAG-hyPBase* and 0.01% Fast Green solution (Sigma-Aldrich, Tokyo, Japan) was injected into a lateral ventricle of each embryo.

### Immunohistochemistry

GRAB_ATP1.0_-expressing mice (4–8 weeks old), which had undergone IUE, were anesthetized with isoflurane (DS Pharma Animal Health Co. Ltd., Osaka, Japan) and transcardially perfused with phosphate-buffered saline (PBS) followed by 4% paraformaldehyde in PBS (Nacalai Tesque, Kyoto, Japan). Brain samples were post-fixed in 4% PFA at 4°C overnight and coronally sectioned at 150 μm thickness using a vibratome-type tissue slicer (DTK-1000, Dosaka-EM, Kyoto, Japan). The brain slices were then immersed in 0.3% Triton X-100 in PBS for 5 min at room temperature. After three time rinses with PBS, the slices were blocked with TNB blocking buffer [0.1 M Tris-HCl, 0.15 M NaCl, 0.5% TSA blocking reagent (PerkinElmer, Boston, MA, U.S.A.)] and incubated with a primary antibody (1:500 in TNB, goat polyclonal anti-GFAP antibody, ab53554, abcam, Cambridge, United Kingdom) and a secondary antibody (1:200 in TNB, Alexa 647 donkey anti-goat IgG, 705-609-147, Jackson ImmunoResearch Laboratories, Inc, West Grove, PA, U.S.A.). The cortical regions were observed using confocal microscopy (FV1000, Olympus, Tokyo, Japan). The mean fluorescence intensity was calculated using Image J (*67*) and transfected and non-transfected hemispheres were compared.

### Cell culture

Primary neuron-glia co-cultures were prepared and cultured as described previously (*37*). In brief, rat primary neuron-glia co-cultures were prepared from 0-day-old (P0) wild-type Sprague-Dawley rat pups (male and female, randomly selected) purchased from Charles River Laboratories (Beijing, China). Hippocampal cells were dissociated from the dissected brains in 0.25% Trypsin-EDTA (Gibco, Waltham, MA, U.S.A.) and plated on 12-mm glass coverslips coated with poly-D-lysine (Sigma-Aldrich) in neurobasal medium (Gibco) containing 2% B-27 supplement (Gibco), 1% GlutaMAX (Gibco), and 1% penicillin-streptomycin (Gibco). Based on glial cell density, after approximately 4 days in culture (DIV 4) cytosine β-D-arabinofuranoside (Sigma-Aldrich) was added to the hippocampal cultures in a 50% growth media exchange, at a final concentration of 2 μM. Primary neuron-glia co-cultures were cultured at 37°C in 5% CO_2_. To express GRAB_ATP1.0_ in neurons, adeno-associated viruses (AAV2/9-hSyn-GRAB_ATP1.0_) were added to neuron-glia co-cultures at DIV 5–9, and DIV ≥13 cells were used for imaging. Before imaging, the culture medium was replaced with Tyrode’s solution containing: 150 mM NaCl, 4 mM KCl, 2 mM MgCl_2_, 2 mM CaCl_2_, 10 mM HEPES, and 10 mM glucose (pH 7.3–7.4). Cells grown on 12-mm coverslips were imaged using a Ti-E A1 confocal microscope (Nikon, Tokyo, Japan) equipped with a 10x/0.45 NA objective, a 20x/0.75 NA objective, a 40x/1.35 NA oil-immersion objective, a 488-nm laser, and a 561-nm laser; green fluorescence (GRAB_ATP1.0_ sensors) and red fluorescence (Calbryte 590, AAT Bioquest) were recorded using a 525/50-nm, and 595/50-nm emission filter, respectively.

### Cranial window surgery

To minimize inflammatory responses, a thin skull surgery with optical clearing methods was employed to prepare a cranial window for *in vivo* imaging (*68*). Mice were anesthetized with the anesthetic agents indicated above, and a metallic head plate (19 mm long, 12 mm wide, and 1 mm thick) with a hole (5 mm in diameter) was attached to the skull with dental cement (GC Corporation, Tokyo, Japan). The skull was thinned with a dental drill and then treated with 10% (W/V) Na_2_EDTA (pH 7.0) for 30–40 min to decalcify the skull. The dip in the skull was filled with 80% glycerol, covered with a circular cover glass (4 mm in diameter, 0.12 mm in thickness, Matsunami, Osaka, Japan), and sealed with cyanoacrylate glue (Aron Alpha, Toagosei CO., LTD., Tokyo, Japan).

### Acute slice preparation

Acute cortical slices were prepared from 4–8 weeks old mice which had undergone IUE. The brain was removed quickly and placed in an ice-cold cutting solution containing: 120 mM Choline-Cl, 3 mM KCl, 1.25 mM NaH_2_PO_4_, 26 mM NaHCO_3_, 8 mM MgCl_2_ and 20 mM D-glucose. Then, the brain was coronally sectioned at 350 μm thickness with a vibratome-type tissue slicer (Pro7, Dosaka-EM, Kyoto, Japan) and incubated for 1 hour at 28°C for recovery in artificial cerebrospinal fluid (ACSF) containing: 124 mM NaCl, 2.5 mM KCl, 1.25 mM NaH_2_PO_4_, 26 mM NaHCO_3_, 2 mM MgCl_2_, 2 mM CaCl_2_, 20 mM D-glucose, and continuously bobbled with 95% O_2_/5% CO_2_. All acute slice experiments were performed within 6 hours after slice recovery to avoid the effects of inflammatory responses (*69*).

### Pharmacological treatments in acute slice experiments

Tetrodotoxin (1 μM; TTX; Latoxan Laboratory, Portes lès Valence, France), DL-Fluorocitrate (100 μM; Sigma-Aldrich, St. Louis, MO, U.S.A.), carbenoxolone (100 μM; Sigma-Aldrich) and bafilomycin A1 (2 μM; L C Laboratories, Woburn, MA, U.S.A.) were each bath applied through superfusion of ACSF during a recording session. DL-Fluorocitrate (100 μM; Sigma-Aldrich) and bafilomycin A1 (2 μM; L C Laboratories), were also pre-incubated for at least 2 hours prior to recording. For calibration of GRAB_ATP1.0_ response in acute slice, ATP was bath-applied at 0.2, 0.5, 1, 5, 10, and 100 μM (Sigma-Aldrich), and astrocytes located at the surface of slices were monitored to avoid the rapid degradation of extracellular ATP by ectonucleotidases deep in the slice.

### Two-photon imaging

Imaging was performed with an in-house two-photon laser scanning microscope controlled by TI workbench software (*70*). GRAB_ATP1.0_ and Lck-REX-GECO1 (*42*) were excited at 920 nm in acute slice experiments and at 860 nm in *in viv*o imaging using a titanium-sapphire pulse laser (Mai Tai DeepSee, Spectra-Physics, Tokyo, Japan) through a 20x objective (XLUMPLFLN20xW, Olympus) in acute slice experiments and through a 25x objective lens (XLPLN25xWMP2, Olympus) in *in viv*o imaging. The emissions were separated by a 580-nm beam splitter, passed through a 495–540 or 573–648-nm band pass filter, and detected with GaAsP-type photomultiplier tubes (H7422PA-40, Hamamatsu Photonics, Hamamatsu, Japan). Time-lapse image sequences were acquired at 1 Hz with a 0.8 × 0.8 μm per pixel resolution, and line-scan images were acquired at 100 Hz. Astrocytes located 30–100 μm below the slice surface were monitored in acute slice experiments; *in vivo*, we imaged astrocytes located less than 140 μm from the brain surface.

### Event detection and overlap analysis in ATP and Ca^2+^ imaging

Background of time-lapse images was determined and subtracted in each image frame by taking the average of >200 pixels of no fluorescence of expressed sensors. Background-subtracted images were temporally smoothed with a three-frame-width moving average filter to reduce noise before event detection analysis in TI Workbench. Additionally, motion artifacts in some data were corrected by ImageJ plug-in StackReg (*71*) implemented in TI Workbench. Processed image data were further analyzed with the Astrocyte Quantitative Analysis (AQuA) software (*72*) in MATLAB for event detection and event feature extraction of ATP and Ca^2+^ events. The overlap between the ATP and Ca^2+^ events was determined by the following criteria (see also Fig. S3): detected areas for ATP and Ca^2+^ events shared pixels; and event durations, defined as within full width at 30% maximum, of both ATP and Ca^2+^ events overlapped with each other. The sequence of the ATP and Ca^2+^ events was then evaluated by the time point at 50% rise, and event pairs that showed a lag of one frame or less were defined as simultaneous events.

### Classification of ATP release events

To classify ATP release events according to the waveform of fluorescent trace, hierarchical clustering was conducted (see also Fig. S5) in Python using NumPy, SciPy, and pandas libraries (*73*–*75*). Firstly, ATP events observed in acute slices, which were not treated with drugs, were collected. Each trace at time points from -40 to 120 frames relative to the 30% rise frame was extracted and scaled with z-score normalization to calculate the shape-based distance. Then, hierarchical clustering was performed based on the shape-based distance using the Ward’s linkage method (*76*). The obtained clusters were used as training samples, and ATP release events observed in pharmacological experiments were classified using the nearest neighbor classification algorithm (*77*). To display the distribution of ATP event waveforms in 2D (Fig. 5G and J), the principal component analysis (PCA) from scikit-learn library (*78*) or the uniform manifold approximation and projection (UMAP) method (*46*) was applied.

### Statistical analysis

Statistics were performed using Python 3.7 and are described in the corresponding figure legend. All tests were two-tailed, and a p-value lower than 0.05 was considered significant. Data following normal distribution were represented as means ± SD with bar plots, and paired t-tests were applied. When normal distribution cannot be assumed, results were represented as the median and interquartile range (IQR) [25th and 75th percentile] with box plots. Mann-Whitney U test and the Wilcoxon signed-rank test were used for a two-group comparison of unpaired and paired data, respectively. In more than three-group comparisons, Steel-Dwass test and Friedman test were used for unpaired and paired data, respectively. The correlation coefficients were calculated using Kendall’s correlation test, a nonparametric method resistant to equal ranking. For evaluating the effect of inhibitors on the frequency of ATP release events assigned to each cluster, p-values were calculated from the Poisson distribution with Bonferroni correction. For assessing the significance of overlapping pattern incidence and Ca^2+^ activity before and after ATP release events, the chance level was calculated by randomly shuffling the event onsets.

## Acknowledgments

YH is supported by the Grant for Doctor 21 from the Yoshida Scholarship Foundation. ZW is supported in part by the Postdoctoral Fellowship of Peking-Tsinghua Center for Life Sciences.

## Funding

This work was supported by Waseda University Grants for Special Research Projects (2019C-715, 2020C-778, 2021C-733, 2022C-629) to TI; IHU FOReSIGHT (ANR-18-IAHU-01) to JL.

## Author contributions

Conceptualization: YH, HF, and TI; investigation: YH and ZW; visualization: YH and ZW; formal analysis: YH and ZW; resources: ZW, TK, JL, YL; supervision: TI; writing—original draft: YH and TI; writing—review & editing: all authors.

## Competing interests

The authors declare that they have no competing interests.

## Data and materials availability

The data supporting the findings of this study are available from the corresponding author upon reasonable request.

## Figures and Tables

**Fig. S1.**
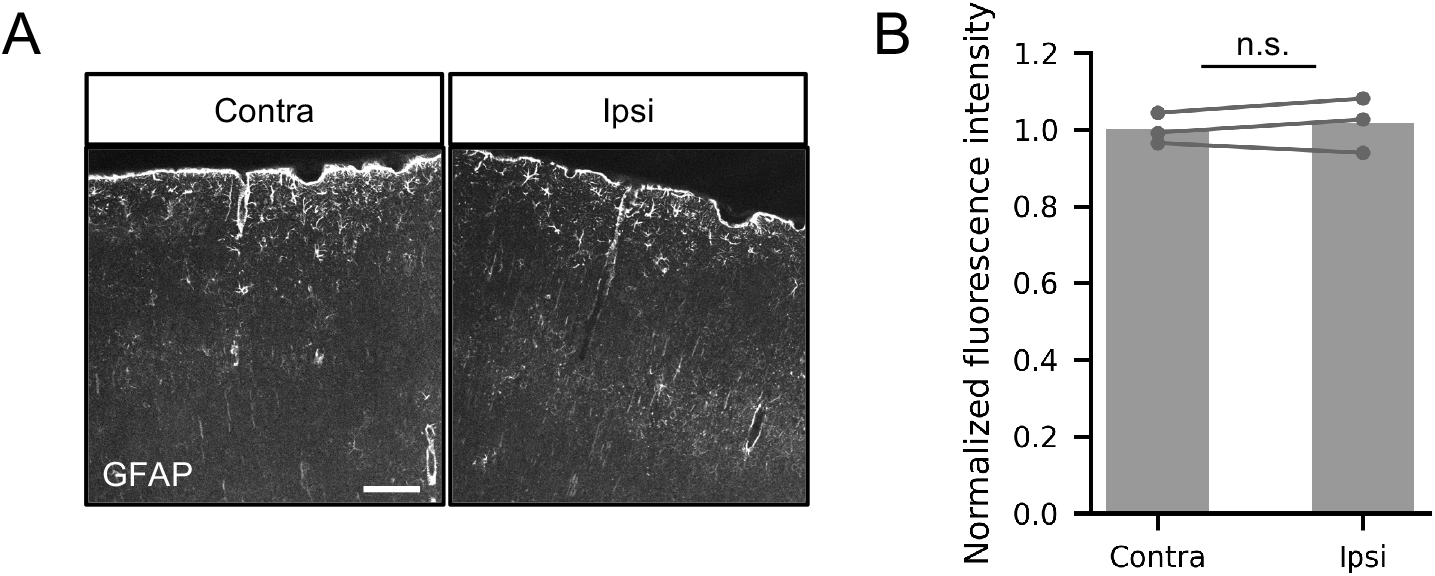
Upregulation of GFAP expression following electroporation is not observed in the cerebral cortex. (**A**) Immunostaining for glial fibrillary acidic protein (GFAP) in the both sides of cortex prepared from a P40 mouse to which *in utero* electroporation had been performed to ipsilateral (Ipsi) side at E15.5. The contralateral (Contra) side was used as a negative control. Scale bar: 100 μm. (**B**) Mean fluorescence intensity in cortex excluding pia mater normalized by the average of control (contralateral side) after subtraction of the background value. Each dot represents an average of six image fields obtained from a mouse. Paired t-test; *p < 0.05; n.s., not significant.

**Fig. S2.**
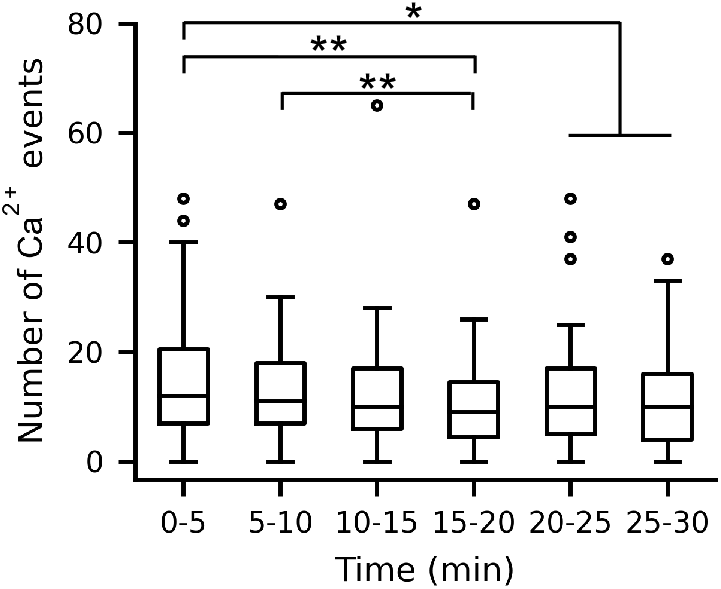
Two-photon imaging does not induce Ca^2+^ hyperactivity in astrocytes. The number of Ca^2+^ events detected per 5 min time window during a 30 min recording session of two-photon imaging in acute slice (n = 55 cells). Friedman test with Bonferroni-corrected Wilcoxon signed-rank test as post-hoc analysis; *p < 0.05; **p < 0.01.

**Fig. S3.**
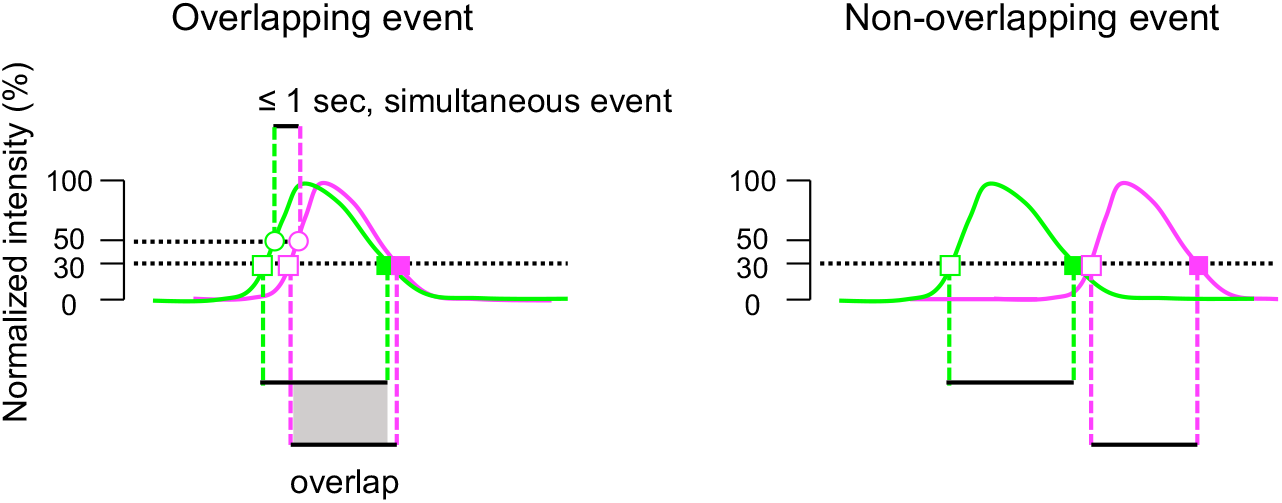
Criteria for overlapping event pair. The overlap between ATP and Ca^2+^ event was judged when the both spatial and temporal criteria meet: areas detected as ATP and Ca^2+^ events by the AQuA software share overlapping portions, and the time course traces overlap each other within the duration of peak width at 30% height of each trace. The order of the overlapping events was then evaluated by the 50% rise time point, and event pairs with the lag within one imaging frame were judged as simultaneous.

**Fig. S4.**
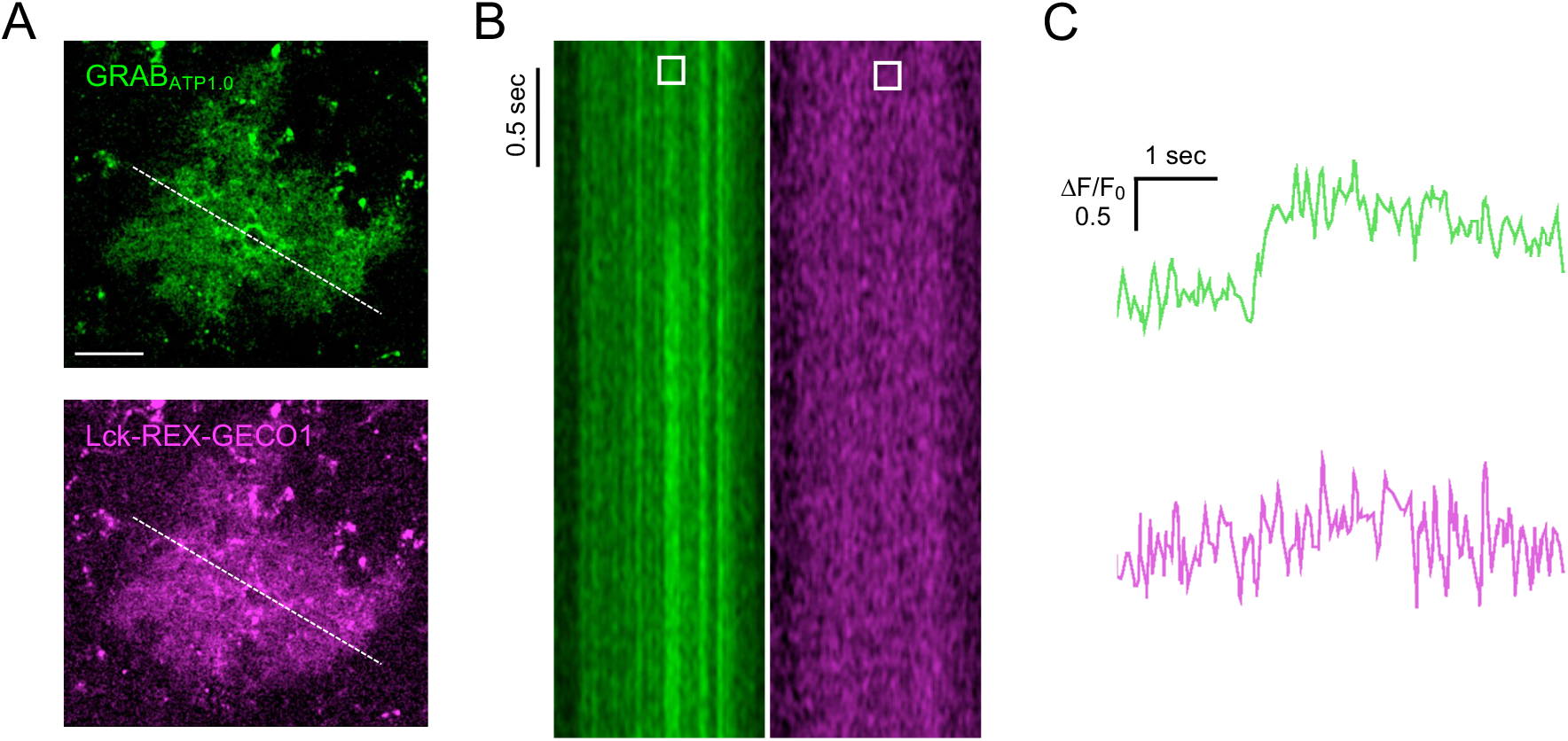
Dual-color imaging with fast line-scanning. (**A**) Images of a cortical astrocyte expressing GRAB_ATP1.0_ (top) and Lck-REX-GECO1 (bottom). White dashed lines represent the location of line-scanning. Scale bars: 100 μm (**B**) Images acquired in the line-scan mode at 100 Hz. (**C**) Time course traces of GRAB_ATP1.0_ and Lck-REX-GECO1 signals obtained from the region indicated with the white rectangle in B. The traces were denoised with a five-frame moving average filter.

**Fig. S5.**
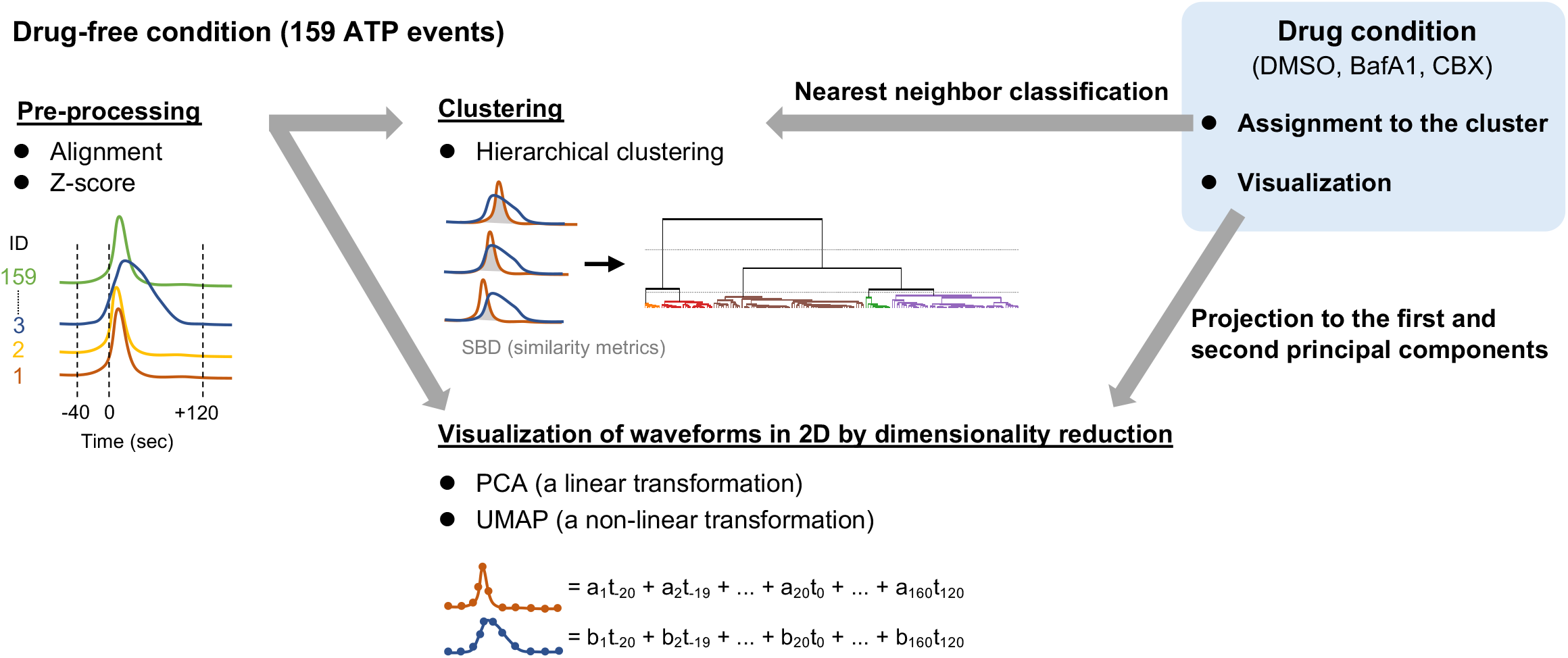
Outline of clustering analysis of ATP release events. ATP release events observed under the drug-free condition in acute slices were used as a training sample for clustering analysis. Events are aligned by the initial rise timing (t0, 30% of the peak amplitude) and 161 time-points from -40 to 120 sec relative to t0 were extracted and scaled with z-score normalization to calculate the shape-based distance (SBD). Then, clusters were determined by hierarchical clustering using SBD and Ward’s linkage method. To visualize the distribution of ATP event waveforms in 2D, the principal component analysis (PCA) or the uniform manifold approximation and projection (UMAP) analysis was applied. ATP release events detected in the pharmacological experiments were assigned to the cluster by calculating SBD and using the nearest neighbor classification algorithm and displayed in 2D by applying PCA with the same parameters used for the drug-free condition.

**Fig. S6.**
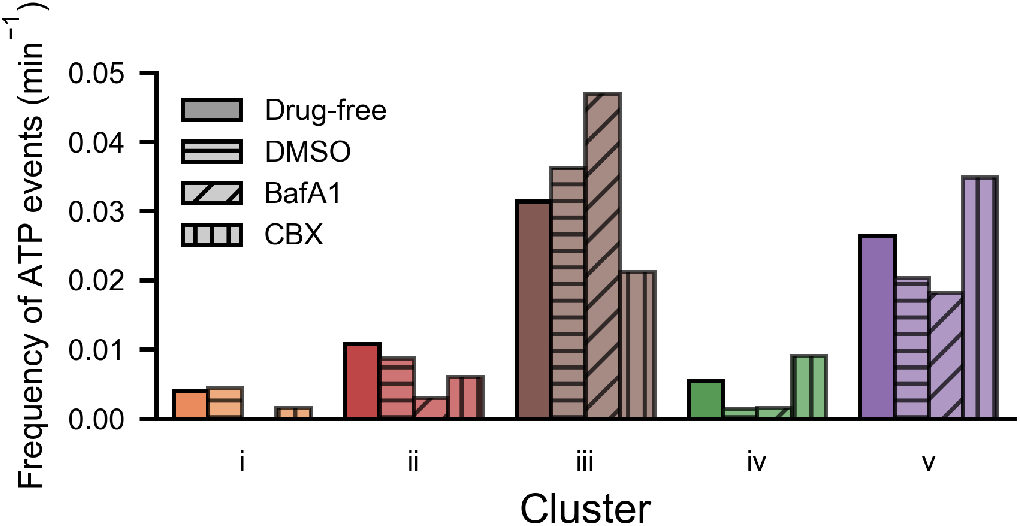
Effects of inhibitors on the ATP release event frequency in the five clusters. Frequency of ATP release events in each of the five clusters in the presence of inhibitors related to Fig. 5.

## Notes

### Competing Interest Statement

The authors have declared no competing interest.

## References

1. M. M. Halassa, P. G. Haydon, Integrated Brain Circuits: Astrocytic Networks Modulate Neuronal Activity and Behavior. Annu. Rev. Physiol. 72, 335–355 (2010).

2. A. Araque, G. Carmignoto, P. G. Haydon, S. H. R. Oliet, R. Robitaille, A. Volterra, Gliotransmitters Travel in Time and Space. Neuron. 81, 728–739 (2014).

3. G. Perea, A. Araque, Astrocytes Potentiate Transmitter Release at Single Hippocampal Synapses. Science. 317, 1083–1086 (2007).

4. C. Henneberger, T. Papouin, S. H. R. Oliet, D. A. Rusakov, Long-term potentiation depends on release of d-serine from astrocytes. Nature. 463, 232–236 (2010).

5. A. Panatier, J. Vallée, M. Haber, K. K. Murai, J.-C. Lacaille, R. Robitaille, Astrocytes Are Endogenous Regulators of Basal Transmission at Central Synapses. Cell. 146, 785–798 (2011).

6. G. Perea, Properties of Synaptically Evoked Astrocyte Calcium Signal Reveal Synaptic Information Processing by Astrocytes. Journal of Neuroscience. 25, 2192–2203 (2005).

7. A. Covelo, A. Araque, Neuronal activity determines distinct gliotransmitter release from a single astrocyte. eLife. 7, e32237 (2018).

8. M. Jun, Q. Xiaolong, Y. Chaojuan, P. Ruiyuan, W. Shukun, W. Junbing, H. Li, C. Hong, C. Jinbo, W. Rong, L. Yajin, M. Lanqun, W. Fengchao, W. Zhiying, A. Jianxiong, W. Yun, Z. Xia, Z. Chen, Y. Zengqiang, Calhm2 governs astrocytic ATP releasing in the development of depression-like behaviors. Mol Psychiatry. 23, 883–891 (2018).

9. X. Cao, L.-P. Li, Q. Wang, Q. Wu, H.-H. Hu, M. Zhang, Y.-Y. Fang, J. Zhang, S.-J. Li, W.-C. Xiong, H.-C. Yan, Y.-B. Gao, J.-H. Liu, X.-W. Li, L.-R. Sun, Y.-N. Zeng, X.-H. Zhu, T.-M. Gao, Astrocyte-derived ATP modulates depressive-like behaviors. Nat Med. 19, 773–777 (2013).

10. Q. Wang, Y. Kong, D.-Y. Wu, J.-H. Liu, W. Jie, Q.-L. You, L. Huang, J. Hu, H.-D. Chu, F. Gao, N.-Y. Hu, Z.-C. Luo, X.-W. Li, S.-J. Li, Z.-F. Wu, Y.-L. Li, J.-M. Yang, T.-M. Gao, Impaired calcium signaling in astrocytes modulates autism spectrum disorder-like behaviors in mice. Nat Commun. 12, 3321 (2021).

11. M. M. Halassa, C. Florian, T. Fellin, J. R. Munoz, S.-Y. Lee, T. Abel, P. G. Haydon, M. G. Frank, Astrocytic Modulation of Sleep Homeostasis and Cognitive Consequences of Sleep Loss. Neuron. 61, 213–219 (2009).

12. S. Koizumi, K. Fujishita, M. Tsuda, Y. Shigemoto-Mogami, K. Inoue, Dynamic inhibition of excitatory synaptic transmission by astrocyte-derived ATP in hippocampal cultures. Proc. Natl. Acad. Sci. U.S.A. 100, 11023–11028 (2003).

13. O. Chever, C.-Y. Lee, N. Rouach, Astroglial Connexin43 Hemichannels Tune Basal Excitatory Synaptic Transmission. Journal of Neuroscience. 34, 11228–11232 (2014).

14. Y. Kamatsuka, M. Fukagawa, T. Furuta, A. Ohishi, K. Nishida, K. Nagasawa, Astrocytes, but Not Neurons, Exhibit Constitutive Activation of P2X7 Receptors in Mouse Acute Cortical Slices under Non-stimulated Resting Conditions. Biological & Pharmaceutical Bulletin. 37, 1958–1962 (2014).

15. S. Chi, Y. Cui, H. Wang, J. Jiang, T. Zhang, S. Sun, Z. Zhou, Y. Zhong, B. Xiao, Astrocytic Piezo1-mediated mechanotransduction determines adult neurogenesis and cognitive functions. Neuron, S0896627322006559 (2022).

16. E. Shigetomi, S. Patel, B. S. Khakh, Probing the Complexities of Astrocyte Calcium Signaling. Trends in Cell Biology. 26, 300–312 (2016).

17. S. Guerra-Gomes, N. Sousa, L. Pinto, J. F. Oliveira, Functional Roles of Astrocyte Calcium Elevations: From Synapses to Behavior. Front. Cell. Neurosci. 11, 427 (2018).

18. A. Semyanov, Spatiotemporal pattern of calcium activity in astrocytic network. Cell Calcium. 78, 15–25 (2019).

19. W. Shen, L. Nikolic, C. Meunier, F. Pfrieger, E. Audinat, An autocrine purinergic signaling controls astrocyte-induced neuronal excitation. Sci Rep. 7, 11280 (2017).

20. S. Mederos, A. Hernández-Vivanco, J. Ramírez-Franco, M. Martín-Fernández, M. Navarrete, A. Yang, E. S. Boyden, G. Perea, Melanopsin for precise optogenetic activation of astrocyte-neuron networks. Glia. 67, 915–934 (2019).

21. Y. Iwai, K. Ozawa, K. Yahagi, T. Mishima, S. Akther, C. T. Vo, A. B. Lee, M. Tanaka, S. Itohara, H. Hirase, Transient Astrocytic Gq Signaling Underlies Remote Memory Enhancement. Front. Neural Circuits. 15, 658343 (2021).

22. N. Takata, H. Hirase, Cortical Layer 1 and Layer 2/3 Astrocytes Exhibit Distinct Calcium Dynamics In Vivo. PLoS ONE. 3, e2525 (2008).

23. M. D. Haustein, S. Kracun, X.-H. Lu, T. Shih, O. Jackson-Weaver, X. Tong, J. Xu, X. W. Yang, T. J. O’Dell, J. S. Marvin, M. H. Ellisman, E. A. Bushong, L. L. Looger, B. S. Khakh, Conditions and Constraints for Astrocyte Calcium Signaling in the Hippocampal Mossy Fiber Pathway. Neuron. 82, 413–429 (2014).

24. R. L. Rungta, L.-P. Bernier, L. Dissing-Olesen, C. J. Groten, J. M. LeDue, R. Ko, S. Drissler, B. A. MacVicar, Ca ^2+^ transients in astrocyte fine processes occur via Ca ^2+^ influx in the adult mouse hippocampus: Ca ^2+^ Transients in Astrocyte Fine Processes. Glia. 64, 2093–2103 (2016).

25. A. Agarwal, P.-H. Wu, E. G. Hughes, M. Fukaya, M. A. Tischfield, A. J. Langseth, D. Wirtz, D. E. Bergles, Transient Opening of the Mitochondrial Permeability Transition Pore Induces Microdomain Calcium Transients in Astrocyte Processes. Neuron. 93, 587-605.e7 (2017).

26. E. Bindocci, I. Savtchouk, N. Liaudet, D. Becker, G. Carriero, A. Volterra, Three-dimensional Ca ^2+^ imaging advances understanding of astrocyte biology. Science. 356, eaai8185 (2017).

27. Y.-W. Wu, S. Gordleeva, X. Tang, P.-Y. Shih, Y. Dembitskaya, A. Semyanov, Morphological profile determines the frequency of spontaneous calcium events in astrocytic processes. Glia. 67, 246–262 (2019).

28. M. Arizono, V. V. G. K. Inavalli, A. Panatier, T. Pfeiffer, J. Angibaud, F. Levet, M. J. T. Ter Veer, J. Stobart, L. Bellocchio, K. Mikoshiba, G. Marsicano, B. Weber, S. H. R. Oliet, U. V. Nägerl, Structural basis of astrocytic Ca2+ signals at tripartite synapses. Nat Commun. 11, 1906 (2020).

29. C. Agulhon, M.-Y. Sun, T. Murphy, T. Myers, K. Lauderdale, T. A. Fiacco, Calcium Signaling and Gliotransmission in Normal vs. Reactive Astrocytes. Front. Pharmacol. 3 (2012), doi:10.3389/fphar.2012.00139.

30. N. Bazargani, D. Attwell, Astrocyte calcium signaling: the third wave. Nat Neurosci. 19, 182–189 (2016).

31. I. Savtchouk, A. Volterra, Gliotransmission: Beyond Black-and-White. J. Neurosci. 38, 14–25 (2018).

32. T. A. Fiacco, K. D. McCarthy, Multiple Lines of Evidence Indicate That Gliotransmission Does Not Occur under Physiological Conditions. J. Neurosci. 38, 3–13 (2018).

33. M. M. Halassa, T. Fellin, H. Takano, J.-H. Dong, P. G. Haydon, Synaptic Islands Defined by the Territory of a Single Astrocyte. Journal of Neuroscience. 27, 6473–6477 (2007).

34. Z. Wu, Y. Li, New frontiers in probing the dynamics of purinergic transmitters in vivo. Neuroscience Research. 152, 35–43 (2020).

35. J. M. Conley, S. Radhakrishnan, S. A. Valentino, M. Tantama, Imaging extracellular ATP with a genetically-encoded, ratiometric fluorescent sensor. PLoS ONE. 12, e0187481 (2017).

36. M. A. Lobas, R. Tao, J. Nagai, M. T. Kronschläger, P. M. Borden, J. S. Marvin, L. L. Looger, B. S. Khakh, A genetically encoded single-wavelength sensor for imaging cytosolic and cell surface ATP. Nat Commun. 10, 711 (2019).

37. Z. Wu, K. He, Y. Chen, H. Li, S. Pan, B. Li, T. Liu, F. Xi, F. Deng, H. Wang, J. Du, M. Jing, Y. Li, A sensitive GRAB sensor for detecting extracellular ATP in vitro and in vivo. Neuron. 110, 770-782.e5 (2022).

38. X. Li, Y. Li, Y. Zhou, J. Wu, Z. Zhao, J. Fan, F. Deng, Z. Wu, G. Xiao, J. He, Y. Zhang, G. Zhang, X. Hu, X. Chen, Y. Zhang, H. Qiao, H. Xie, Y. Li, H. Wang, L. Fang, Q. Dai, Real-time denoising enables high-sensitivity fluorescence time-lapse imaging beyond the shot-noise limit. Nat Biotechnol (2022), doi:10.1038/s41587-022-01450-8.

39. T. Kumamoto, F. Maurinot, R. Barry-Martinet, C. Vaslin, S. Vandormael-Pournin, M. Le, M. Lerat, D. Niculescu, M. Cohen-Tannoudji, A. Rebsam, K. Loulier, S. Nedelec, S. Tozer, J. Livet, Direct Readout of Neural Stem Cell Transgenesis with an Integration-Coupled Gene Expression Switch. Neuron. 107, 617-630.e6 (2020).

40. B. S. Khakh, R. A. North, Neuromodulation by Extracellular ATP and P2X Receptors in the CNS. Neuron. 76, 51–69 (2012).

41. L. Vitiello, S. Gorini, G. Rosano, A. la Sala, Immunoregulation through extracellular nucleotides. Blood. 120, 511–518 (2012).

42. J. Wu, A. S. Abdelfattah, L. S. Miraucourt, E. Kutsarova, A. Ruangkittisakul, H. Zhou, K. Ballanyi, G. Wicks, M. Drobizhev, A. Rebane, E. S. Ruthazer, R. E. Campbell, A long Stokes shift red fluorescent Ca2+ indicator protein for two-photon and ratiometric imaging. Nat Commun. 5, 5262 (2014).

43. E. Schmidt, M. Oheim, Infrared Excitation Induces Heating and Calcium Microdomain Hyperactivity in Cortical Astrocytes. Biophysical Journal. 119, 2153–2165 (2020).

44. P. Illes, G. Burnstock, Y. Tang, Astroglia-Derived ATP Modulates CNS Neuronal Circuits. Trends in Neurosciences. 42, 885–898 (2019).

45. J. Paparrizos, L. Gravano, “k-Shape: Efficient and Accurate Clustering of Time Series” in Proceedings of the 2015 ACM SIGMOD International Conference on Management of Data (ACM, Melbourne Victoria Australia, 2015; https://dl.acm.org/doi/10.1145/2723372.2737793), pp. 1855–1870.

46. L. McInnes, J. Healy, J. Melville, UMAP: Uniform Manifold Approximation and Projection for Dimension Reduction. arXiv (2018), doi:10.48550/ARXIV.1802.03426.

47. T. Kikuchi, J. Gonzalez-Soriano, A. Kastanauskaite, R. Benavides-Piccione, A. Merchan-Perez, J. DeFelipe, L. Blazquez-Llorca, Volume Electron Microscopy Study of the Relationship Between Synapses and Astrocytes in the Developing Rat Somatosensory Cortex. Cerebral Cortex. 30, 3800–3819 (2020).

48. P. B. Guthrie, J. Knappenberger, M. Segal, M. V. L. Bennett, A. C. Charles, S. B. Kater, ATP Released from Astrocytes Mediates Glial Calcium Waves. J. Neurosci. 19, 520–528 (1999).

49. A. Delekate, M. Füchtemeier, T. Schumacher, C. Ulbrich, M. Foddis, G. C. Petzold, Metabotropic P2Y1 receptor signalling mediates astrocytic hyperactivity in vivo in an Alzheimer’s disease mouse model. Nat Commun. 5, 5422 (2014).

50. S. Aten, C. M. Kiyoshi, E. P. Arzola, J. A. Patterson, A. T. Taylor, Y. Du, A. M. Guiher, M. Philip, E. G. Camacho, D. Mediratta, K. Collins, K. Boni, S. A. Garcia, R. Kumar, A. N. Drake, A. Hegazi, L. Trank, E. Benson, G. Kidd, D. Terman, M. Zhou, Ultrastructural view of astrocyte arborization, astrocyte-astrocyte and astrocyte-synapse contacts, intracellular vesicle-like structures, and mitochondrial network. Progress in Neurobiology. 213, 102264 (2022).

51. D. Li, K. Hérault, K. Zylbersztejn, M. A. Lauterbach, M. Guillon, M. Oheim, N. Ropert, Astrocyte VAMP3 vesicles undergo Ca ^2+^ -independent cycling and modulate glutamate transporter trafficking: Trafficking of astrocyte VAMP3 vesicles. J Physiol. 593, 2807–2832 (2015).

52. C. Giaume, L. Leybaert, C. C. Naus, J. C. Sáez, Connexin and pannexin hemichannels in brain glial cells: properties, pharmacology, and roles. Front. Pharmacol. 4 (2013), doi:10.3389/fphar.2013.00088.

53. D. A. Sahlender, I. Savtchouk, A. Volterra, What do we know about gliotransmitter release from astrocytes? Phil. Trans. R. Soc. B. 369, 20130592 (2014).

54. J. Stehberg, R. Moraga-Amaro, C. Salazar, A. Becerra, C. Echeverría, J. A. Orellana, G. Bultynck, R. Ponsaerts, L. Leybaert, F. Simon, J. C. Sáez, M. A. Retamal, Release of gliotransmitters through astroglial connexin 43 hemichannels is necessary for fear memory consolidation in the basolateral amygdala. FASEB j. 26, 3649–3657 (2012).

55. N. Prochnow, A. Abdulazim, S. Kurtenbach, V. Wildförster, G. Dvoriantchikova, J. Hanske, E. Petrasch-Parwez, V. I. Shestopalov, R. Dermietzel, D. Manahan-Vaughan, G. Zoidl, Pannexin1 Stabilizes Synaptic Plasticity and Is Needed for Learning. PLoS ONE. 7, e51767 (2012).

56. S. O. Suadicani, R. Iglesias, J. Wang, G. Dahl, D. C. Spray, E. Scemes, ATP signaling is deficient in cultured pannexin1-null mouse astrocytes. Glia. 60, 1106–1116 (2012).

57. Y. Fujii, S. Maekawa, M. Morita, Astrocyte calcium waves propagate proximally by gap junction and distally by extracellular diffusion of ATP released from volume-regulated anion channels. Sci Rep. 7, 13115 (2017).

58. H.-T. Liu, R. Z. Sabirov, Y. Okada, Oxygen-glucose deprivation induces ATP release via maxi-anion channels in astrocytes. Purinergic Signalling. 4, 147–154 (2008).

59. A. Taruno, ATP Release Channels. IJMS. 19, 808 (2018).

60. J. Zhang, H. Wang, C. Ye, W. Ge, Y. Chen, Z. Jiang, C. Wu, M. Poo, S. Duan, ATP Released by Astrocytes Mediates Glutamatergic Activity-Dependent Heterosynaptic Suppression. Neuron. 40, 971–982 (2003).

61. Z. Tan, Y. Liu, W. Xi, H. Lou, L. Zhu, Z. Guo, L. Mei, S. Duan, Glia-derived ATP inversely regulates excitability of pyramidal and CCK-positive neurons. Nat Commun. 8, 13772 (2017).

62. U. Lalo, O. Palygin, S. Rasooli-Nejad, J. Andrew, P. G. Haydon, Y. Pankratov, Exocytosis of ATP From Astrocytes Modulates Phasic and Tonic Inhibition in the Neocortex. PLoS Biol. 12, e1001747 (2014).

63. S. E. Haynes, G. Hollopeter, G. Yang, D. Kurpius, M. E. Dailey, W.-B. Gan, D. Julius, The P2Y12 receptor regulates microglial activation by extracellular nucleotides. Nat Neurosci. 9, 1512–1519 (2006).

64. E. Császár, N. Lénárt, C. Cserép, Z. Környei, R. Fekete, B. Pósfai, D. Balázsfi, B. Hangya, A. D. Schwarcz, E. Szabadits, D. Szöllősi, K. Szigeti, D. Máthé, B. L. West, K. Sviatkó, A. R. Brás, J.-C. Mariani, A. Kliewer, Z. Lenkei, L. Hricisák, Z. Benyó, M. Baranyi, B. Sperlágh, Á. Menyhárt, E. Farkas, Á. Dénes, Microglia modulate blood flow, neurovascular coupling, and hypoperfusion via purinergic actions. Journal of Experimental Medicine. 219, e20211071 (2022).

65. K. Motohashi, A simple and efficient seamless DNA cloning method using SLiCE from Escherichia coli laboratory strains and its application to SLiP site-directed mutagenesis. BMC Biotechnol. 15, 47 (2015).

66. B. Li, M. Chavarha, Y. Kobayashi, S. Yoshinaga, K. Nakajima, M. Z. Lin, T. Inoue, Two-Photon Voltage Imaging of Spontaneous Activity from Multiple Neurons Reveals Network Activity in Brain Tissue. iScience. 23, 101363 (2020).

67. C. A. Schneider, W. S. Rasband, K. W. Eliceiri, NIH Image to ImageJ: 25 years of image analysis. Nat Methods. 9, 671–675 (2012).

68. Y.-J. Zhao, T.-T. Yu, C. Zhang, Z. Li, Q.-M. Luo, T.-H. Xu, D. Zhu, Skull optical clearing window for in vivo imaging of the mouse cortex at synaptic resolution. Light Sci Appl. 7, 17153–17153 (2018).

69. H. Chai, B. Diaz-Castro, E. Shigetomi, E. Monte, J. C. Octeau, X. Yu, W. Cohn, P. S. Rajendran, T. M. Vondriska, J. P. Whitelegge, G. Coppola, B. S. Khakh, Neural Circuit-Specialized Astrocytes: Transcriptomic, Proteomic, Morphological, and Functional Evidence. Neuron. 95, 531-549.e9 (2017).

70. T. Inoue, TI Workbench, an integrated software package for electrophysiology and imaging. Microscopy. 67, 129–143 (2018).

71. P. Thévenaz, U. E. Ruttimann, M. Unser, A pyramid approach to subpixel registration based on intensity. IEEE Trans. on Image Process. 7, 27–41 (1998).

72. Y. Wang, N. V. DelRosso, T. V. Vaidyanathan, M. K. Cahill, M. E. Reitman, S. Pittolo, X. Mi, G. Yu, K. E. Poskanzer, Accurate quantification of astrocyte and neurotransmitter fluorescence dynamics for single-cell and population-level physiology. Nat Neurosci. 22, 1936–1944 (2019).

73. C. R. Harris, K. J. Millman, S. J. van der Walt, R. Gommers, P. Virtanen, D. Cournapeau, E. Wieser, J. Taylor, S. Berg, N. J. Smith, R. Kern, M. Picus, S. Hoyer, M. H. van Kerkwijk, M. Brett, A. Haldane, J. F. Del Río, M. Wiebe, P. Peterson, P. Gérard-Marchant, K. Sheppard, T. Reddy, W. Weckesser, H. Abbasi, C. Gohlke, T. E. Oliphant, Array programming with NumPy. Nature. 585, 357–362 (2020).

74. W. McKinney, Data Structures for Statistical Computing in Python. Proceedings of the 9th Python in Science Conference. 445, 51–56 (2010).

75. P. Virtanen, R. Gommers, T. E. Oliphant, M. Haberland, T. Reddy, D. Cournapeau, E. Burovski, P. Peterson, W. Weckesser, J. Bright, S. J. van der Walt, M. Brett, J. Wilson, K. J. Millman, N. Mayorov, A. R. J. Nelson, E. Jones, R. Kern, E. Larson, C. J. Carey, İ. Polat, Y. Feng, E. W. Moore, J. VanderPlas, D. Laxalde, J. Perktold, R. Cimrman, I. Henriksen, E. A. Quintero, C. R. Harris, A. M. Archibald, A. H. Ribeiro, F. Pedregosa, P. van Mulbregt, SciPy 1.0 Contributors, SciPy 1.0: fundamental algorithms for scientific computing in Python. Nat Methods. 17, 261–272 (2020).

76. J. H. Ward, Hierarchical Grouping to Optimize an Objective Function. Journal of the American Statistical Association. 58, 236–244 (1963).

77. E. Fixt, J. L. Hodges, Discriminatory Analysis. Nonparametric Discrimination: Consistency Properties. US Air Force School of Aviation Medicine. Technical Report 4, 11 (1951).

78. F. Pedregosa, G. Varoquaux, A. Gramfort, V. Michel, B. Thirion, O. Grisel, M. Blondel, P. Prettenhofer, R. Weiss, V. Dubourg, J. Vanderplas, A. Passos, D. Cournapeau, Scikit-learn: Machine Learning in Python. Journal of Machine Learning Research. 12, 2825–2830 (2011).

